# A reproducible and tunable synthetic soil microbial community provides new insights into microbial ecology

**DOI:** 10.1101/2022.05.19.492707

**Authors:** Joanna Coker, Kateryna Zhalnina, Clarisse Marotz, Deepan Thiruppathy, Megan Tjuanta, Gavin D’Elia, Rodas Hailu, Talon Mahosky, Meagan Rowan, Trent R. Northen, Karsten Zengler

## Abstract

Microbial soil communities form commensal relationships with plants to promote the growth of both parties. Optimization of plant-microbe interactions to advance sustainable agriculture is an important field in agricultural research. However, investigation in this field is hindered by a lack of model microbial community systems and efficient approaches for building these communities. Two key challenges in developing standardized model communities are maintaining community diversity over time and storing/resuscitating these communities after cryopreservation, especially considering the different growth rates of organisms. Here, a model community of 17 soil microorganisms commonly found in the rhizosphere of diverse plant species, isolated from soil surrounding a single switchgrass plant, has been developed and optimized for use with fabricated ecosystem devices (EcoFABs). EcoFABs allow reproducible research in model plant systems, with precise control of environmental conditions and easy measurement of plant-microbe metrics. The model soil community grows reproducibly *in vitro* between replicates and experiments, with high community α-diversity achieved through growth in low-nutrient media and adjustment of starting composition ratios for the growth of individual organisms. The community additionally grows in EcoFAB devices and regrows with a similar composition to unfrozen communities following cryopreservation with glycerol, allowing for dissemination of the model community. Our results demonstrate the generation of a stable microbial community that can be used with EcoFAB devices and shared between research groups for maximum reproducibility.

**Importance:** Microbes associate with plants in distinct soil communities, to the benefit of both the soil microbes and the plant. Interactions between plants and these microbes can improve plant growth and health and are therefore a field of study in sustainable agricultural research. In this study, a model community of 17 soil bacteria has been developed to further reproducible study of plant-soil microbe interactions. Preservation of the microbial community has been optimized for dissemination to other research settings. Overall, this work will advance soil microbe research through optimization of a robust, reproducible model community.

## Introduction

The scientific community has developed robust model systems for research in animals, plants, and individual microbes^1,2^. These systems allow experiments to be repeated and validated across research groups, leading to a body of research that grows on the work of others. However, microbiome research currently lacks widely-accepted reproducible model systems, despite the recognition that microbial communities play a fundamental role in biological systems^3–5^. Indeed, host organisms and their microbiota are often referred to as one meta-organism, requiring both parts of the system to thrive^6–8^. Several groups have worked to develop reproducible microbial systems, such as a microbial chemostat^9^; the Lubbock chronic-wound biofilm model^10^; or *in vitro* gut systems incorporating microbes^11–13^, most notably the Altered Schaedler Flora community^14,15^. These systems address important research questions about the interactions between microbes and their host environment. However, they normally do not probe the mechanisms of host-community interactions, particularly in plant-microbe communities under environmental perturbations.

To address specific questions pertaining to the inner workings of microbial communities, researchers must be able to alter the presence and abundance of specific organisms, introduce genetic alterations as necessary, and maintain strict control of growth conditions like temperature, humidity, acidity, and light^8,16–18^. At present, experiments with synthetic microbial communities are the only viable method to design research studies within these constraints^8,19–21^. Although the use of bioengineering tools to introduce specific changes in natural microbial communities shows promise in this area^22–24^, these systems still lack an ability to predict the effect of engineering outcomes on the community as a whole^24^. Synthetic microbial communities therefore represent the most suitable approach to investigate community dynamics.

Plant-microbiome interactions have been the focus of an increasing number of studies in recent years, especially with the potential to optimize agricultural production through the promotion of plant growth and soil health^5,8,25^. These studies clearly show that plant microbiome communities are heavily influenced by the location of microbial colonization on the plant^21,26,27^ and the host plant genotype^28,29^. Each of these studies, some explicitly and some implicitly, are searching for what has been termed the “minimal microbial community”, the minimal set of organisms required to accurately reproduce natural community functions^16^. The number of community microbes in studies ranged from under 10^21,28^ to between 20 and 40^27,29–31^, although some studies starting with a high number of microbes reported that only a small number of organisms consistently colonized plant sites^21,30^. In addition to loss of starting organisms, *in vitro* microbial communities commonly lose α-diversity over time, compared to the starting community^32–34^. The vast majority of synthetic microbial communities are constructed with equal amounts of each organism, although Bai et al.^27^ compared an equal (1:1:1:1) ratio of four represented phyla to an unequal ratio (1:1:1:0.25) but found the final community compositions were similar. However, it has recently been shown that starting ratios, even in a simple co-culture, can have a significant effect on community growth and composition^35,36^. To what extent equal ratios in the starting inoculum produce the most reproducible and diverse synthetic community is still an open question^36^. When generating soil communities, we hypothesized that synthetic community α-diversity could be increased by adjustment of starting organism ratios, with higher levels of organisms that decrease in abundance during growth of an equally-mixed community.

Here, we present the generation of a diverse, reproducible, and tunable synthetic microbial community, composed of soil bacteria isolates obtained from switchgrass agricultural fields. Using a picoliter liquid printer to allow precise control of the initial bacterial inoculum, we tested over 20 community starting composition ratios to generate a synthetic community with maximum robustness and α-diversity. We then used this community to probe the effect of DNA from dead cells on sequencing composition results. To further support the reproducibility of this model community, we additionally determined a method for the cryopreservation of the community enabling it to be shared with other research groups. The 17-member community can readily be applied to EcoFAB devices, which allow reproducible research in model plant systems with precise control of environmental conditions and easy measurement of plant-microbe metrics^18,37^.

## Results

### Strain selection

Our overall goal was to generate a stable, reproducible microbial community for use with EcoFAB devices to study plant-microbe interactions in the rhizosphere. To this end, we selected 18 microbial strains isolated from the rhizosphere and bulk soil surrounding a single switchgrass plant that span the typical diversity found in the rhizosphere of grasses or food crops (Supplemental Table 1). One strain was later eliminated to result in a final 17-member community, described below. These strains were all from different genera, to facilitate community diversity and ease of strain identification through 16S rRNA gene sequencing in the final community. All strains can be grown axenically *in vitro* under aerobic conditions without shaking in liquid Reasoner’s 2A (R2A) media at 30 °C (see Materials and Methods).

We assembled these strains into synthetic communities using a SCIENION CellenONE liquid-handling robot printer (SCIENION US Inc., Phoenix, AZ) (Figure 1). The CellenONE machine is capable of dispensing droplets from 300-450 picoliters. Liquid samples can be taken up into the piezo dispense capillary (PDC), then dispensed in individual drops of precise volume through a piezoelectric pulse. The machine can be programmed to dispense drops from the PDC in specific locations or patterns on a target of the operator’s choosing. For the soil communities, individual strains in diluted liquid culture were taken up from wells of a 96- or 384-well plate and dispensed to a target 96-well plate with fresh liquid growth media. However, samples can also be dispensed to targets such as 384-well plates, microscope slides, or agar plates. Use of this system allowed the 18 strains to be combined in varying starting ratios by programming a different number of starting drops per strain for each community. Individual strains were diluted to an optical density at 600nm (OD_600_) of 0.025 for each experiment.

**Figure 1.**
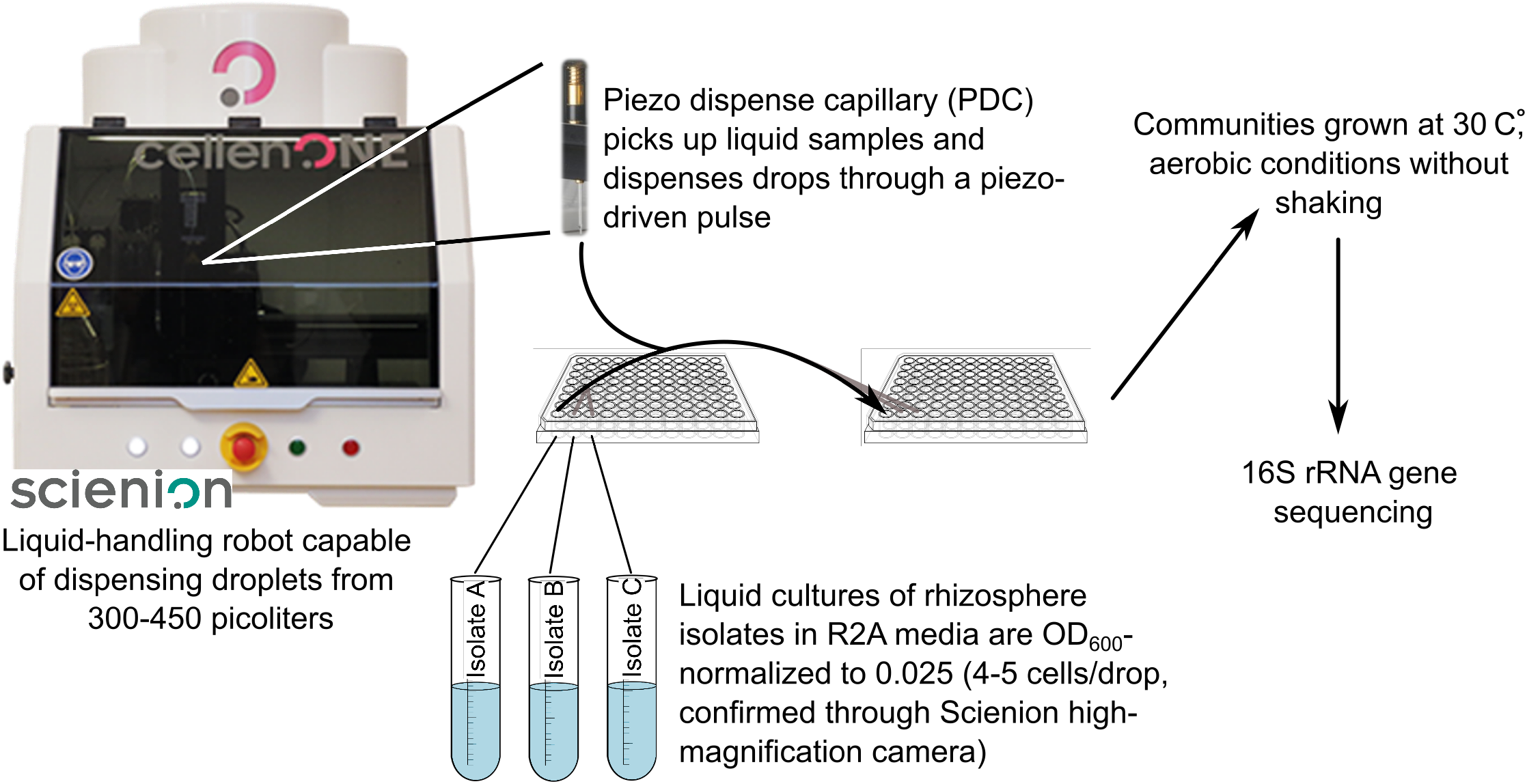
Schematic of synthetic rhizosphere community generation using a piezo dispense capillary (PDC) device. Isolates were grown for 3-4 days in liquid R2A media, then OD_600_-normalized to 0.025 and loaded into individual wells in the probe plate. The PDC drew liquid up from one well of the probe plate and dispensed a programmed number of drops in desired wells of the target plate. This process was repeated for each isolate to result in a final mixed community. Communities were grown aerobically for the desired amount of time, then analyzed for composition and diversity with 16S rRNA sequencing.

Following community assembly, communities were allowed to grow for up to 11 days. Growth was monitored through OD_600_. Community growth was halted at the desired time point by freezing the communities at −20 °C. DNA was extracted from communities by heating samples to 95 °C in a PCR machine for 10 minutes. 5μl of undiluted supernatant from heated community samples was used to generate 16S rRNA gene sequencing libraries. This method was confirmed to produce the same 16S sequencing results as DNA extraction with a commercial DNA extraction kit (Qiagen PowerSoil Pro) (Supplemental Figure 1A-B).

### Automated assembly of synthetic communities produces similar results to hand assembly

The use of the picoliter printer to assemble synthetic communities was chosen to increase throughput and to potentially reduce variability from human and calibration error during pipetting. We therefore compared the diversity and composition of eight replicates of an automated-assembly community (machine) to sixteen replicates of a hand-assembly community (human) after 3 days of growth in 0.1X R2A media. The hand-assembled communities were composed of 4-6 replicates each from 4 different lab members. The growth rate and final OD_600_ value was the same between machine- and hand-assembled communities (Figure 2A). The β-diversity metric of Bray-Curtis distance showed a significantly larger dissimilarity between communities assembled by hand compared to machine for two of the four people (one-way ANOVA with Benjamini-Hochberg FDR, * p<0.05 *** p<0.001) (Figure 2B). Similarly, α-diversity showed a greater spread in hand-assembled than machine-assembled communities for two of the four people (Supp Figure 1C). These results indicate that community assembly with the automated printer will, on average, result in less variability than community assembly by different people.

**Figure 2.**
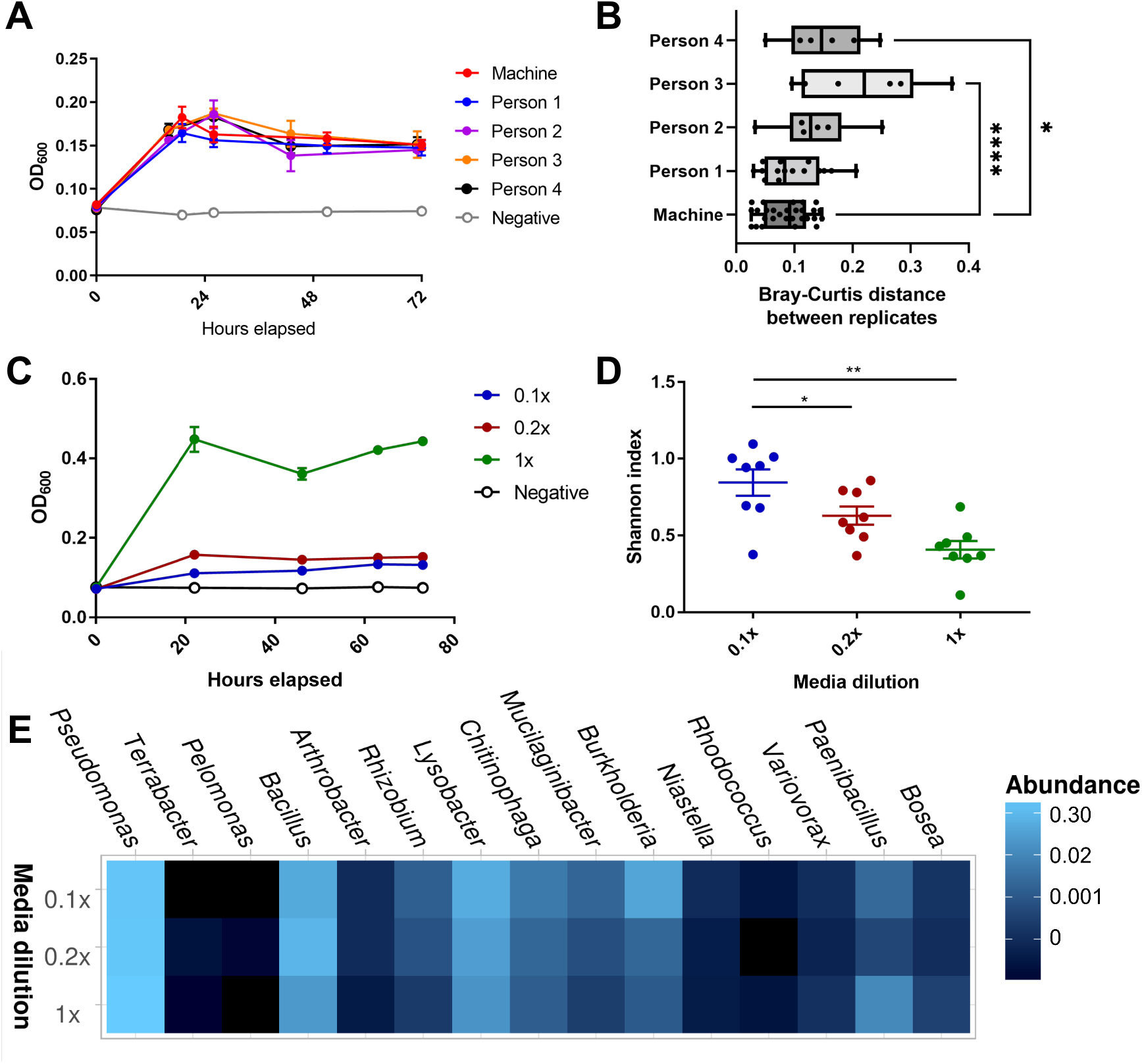
Community diversity with hand-assembly and media dilutions. **A)** OD_600_ values of human-assembled (human) and machine-assembled communities (machine) over 3 days (72 hours) of growth (n=6-8 each). **B)** Bray-Curtis distance on 16S rRNA gene amplicon sequencing between replicates of the human- and machine-assembled communities. **C)** OD_600_ values of machine-assembled equally-mixed communities in 1X, 0.2X, and 0.1X R2A media over 3 days (n=8 each). **D)** Observed OTUs (left) and Shannon diversity index (right) of media dilution communities (Student’s t-test, * = p<0.05, **** = p<0.001). **E)** Heatmap of taxonomy relative abundance of media dilution communities from 16S sequencing. Taxonomic order was determined by PCoA clustering of the Bray-Curtis distance. Replicates for each condition were merged with the Phyloseq command merge_samples(group = “Media_dilution”).

### α-diversity of the synthetic soil community is enhanced through low-nutrient conditions

We next sought to test if nutrient availability affected the growth of individual strains within the community. We compared the growth and diversity of an equally-mixed community of all 18 strains between 1X, 0.2X, and 0.1X R2A media (n=8 for each condition) after 3 days of growth. As expected, total community growth was highest in 1X media, followed by 0.2X and 0.1X media (Figure 2C). However, the α-diversity metrics of observed OTUs and Shannon diversity index were lowest in 1X media and increased as media dilution increased (Figure 2D, Supp Figure 1D). Taxonomy analysis of 16S sequencing data revealed that the *Pseudomonas* strain commonly grew to a high final proportion of the final community, regardless of media dilution (Figure 2E). However, the 0.1X communities displayed higher relative abundances of other, less-dominant strains, such as *Chitinophaga, Burkholderia*, and *Mucilaginibacter*. Individual growth curves of all organisms can be found in Supplemental Figure 2.

### α-diversity of the synthetic soil community is maximized through adjustment of starting community ratios

We next sought to maximize community diversity through adjustment of community starting ratios, meaning organisms were mixed in the starting community in ratios other than 1:1. We tested 11 different starting ratios with and without *Pseudomonas* (22 ratios total; Figure 3A). The exact calculations and ratios are provided in Supplemental Tables 1-2. Briefly, the ratios were calculated based on the change in relative abundance after 3 days of growth from an equally-mixed inoculum. The starting relative abundance (SRA; relative abundance in the inoculum), final relative abundance (FRA; relative abundance after 3 days of growth), and FRA/SRA ratio (FSR) values were applied with various equations to try to design communities with high α-diversity. When designing the community compositions, we hypothesized that starting with lower amounts of organisms with a high SRA or FSR and higher amounts of organisms with a low SRA or FSR would increase α-diversity (Figure 3A, “FSR-based” and “SRA-based” compositions). We also included 4 compositions with different starting amounts of an equally-mixed community (Figure 3A, “Equal” compositions). For this study, two identical 96-well plates were assembled at the same time with the picoliter printer and allowed to grow for 2 and 6 days, respectively.

**Figure 3.**
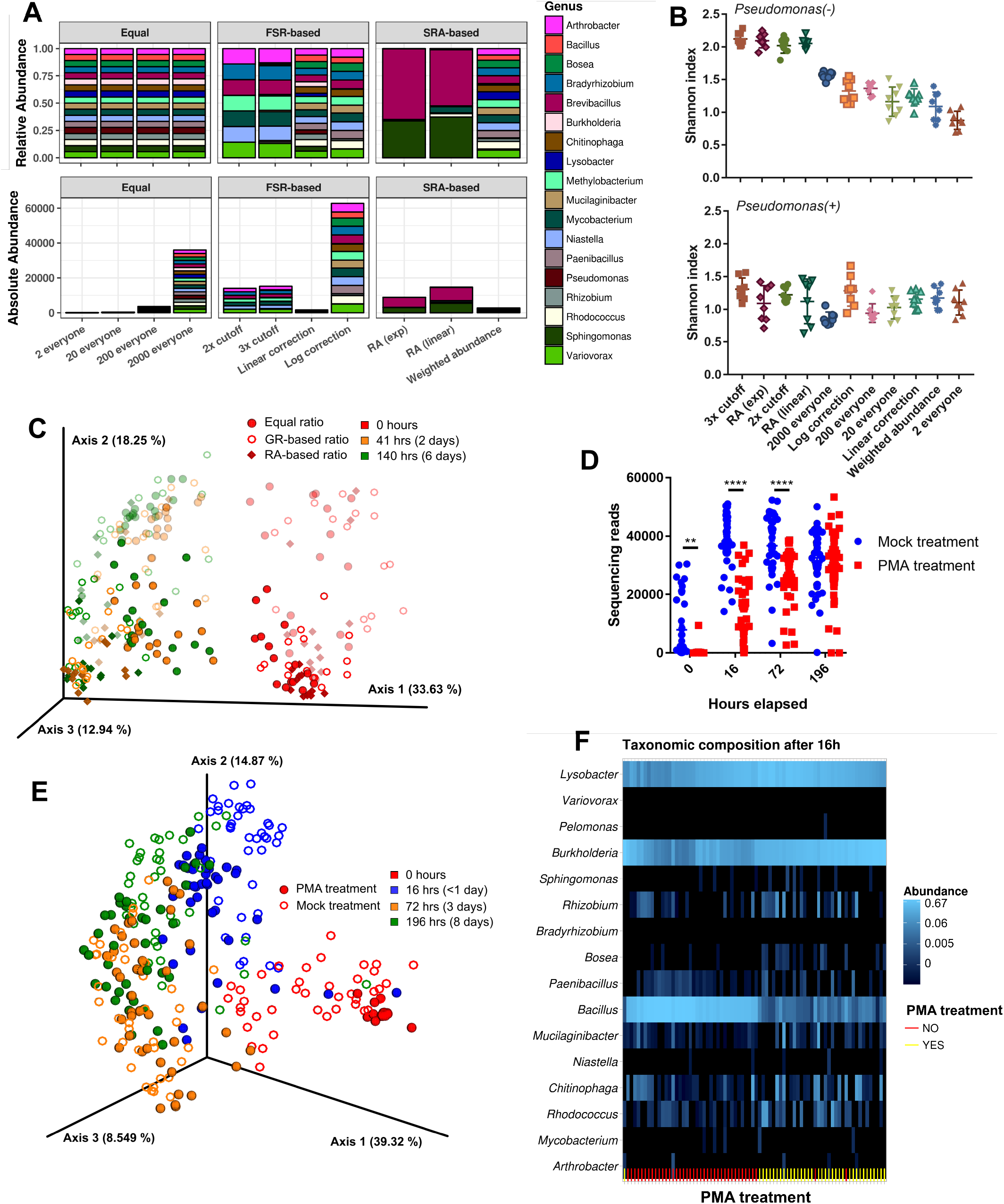
Community diversity with starting ratio adjustments and removal of relic DNA. A) Representation of the 11 community starting ratios used in this study, as both relative (top) and absolute (bottom) abundances. Descriptive names of the ratios are on the x-axis. For equal communities, all organisms were added in equal but increasing amounts; the number refers to the number of drops released by the printer for each organism. For FRA-based and SRA-based adjusted communities, the number of drops for each organism was calculated as shown in Supplemental Tables 2-3. For communities without *Pseudomonas, Pseudomonas* was not added. **B)** Shannon diversity index of each community ratio, 2 and 6 days combined. Communities without *Pseudomonas* are on top, communities with *Pseudomonas* on bottom. Communities are shown in order of decreasing average Shannon index for communities without *Pseudomonas*. (n = 4 each) **C)** PCA of robust Aitchison distance between communities with different starting ratios. Symbols of communities with *Pseudomonas* have reduced opacity. GR = growth rate; RA = relative abundance. **D)** Number of sequencing reads passing quality filtering per sample for PMA- and mock-treatment conditions. **E)** PCA of Aitchison distance between PMA- and mock-treatment communities. **F)** Heatmap of taxonomic composition of the 5 community ratios after 16 hours of growth. Sample order on the x-axis was determined by hierarchical clustering of Bray-Curtis distance in the Phyloseq package. PMA-treated (yellow) and mock-treated (red) communities are marked in the rug plot at the bottom of each heatmap.

In general, communities containing *Pseudomonas* grew to slightly higher OD_600_ values than communities without *Pseudomonas* (Supplemental Figure 3). α-diversity, as measured by Shannon index, was highest in the following communities without *Pseudomonas*: 2x cutoff, in which organisms with FSR<0.05 received 2000 drops from the starting isolate culture while organisms with FSR>0.5 received 2 drops; 3x cutoff, in which organisms with FSR>1, FSR 0.05-1, and FSR<0.05 received 2000, 200, and 2 drops respectively; RA (exp), in which the number of drops decreases exponentially with FRA; and RA (linear), in which the number of drops decreases linearly with FRA (Figure 3B). Analysis of robust Aitchison distance, as a metric of β-diversity, showed that 2- and 6-day communities were significantly different from starting communities (Figure 3C; pairwise PERMANOVA with Benjamini-Hochberg FDR correction, p = 0.0015). Communities also separated between those with and without *Pseudomonas* (pairwise PERMANOVA with Benjamini-Hochberg FDR correction, p = 0.001)

A well-known issue with genomics analysis is the inability to distinguish DNA from dead cells or other extracellular sources (“relic DNA”) from live cell DNA after sequencing^38–40^. To determine the effect of relic DNA on our synthetic community samples, we compared untreated communities to communities treated with propidium monoazide (PMA) to remove extracellular DNA prior to sequencing^40^. The communities with the five highest α-diversity values from Figure 3B were chosen to examine the effect of relic DNA. Four identical plates were prepared with the picoliter printer, with plates collected as time points at 0, 16, 72, and 196 hours. Overall community growth was not significantly different between communities (Supplemental Figure 4A-B).

16S rRNA gene sequencing of PMA-treated communities showed significantly fewer reads passing quality filtration at the 0-, 16-, and 72-hour timepoints compared to mock-treatment communities, although the gap decreased as time increased (Figure 4D). No difference was seen in the number of reads between mock- and PMA-treated communities at 196 hours. PCA of Aitchison distance between the communities showed a separation between the 0-hour and other timepoints (Figure 4E). PMA-treated samples were significantly different from the mock-treated samples at 0, 16, and 196 hours, but not at the 72-hour time point (pairwise PERMANOVA with Benjamini-Hochberg FDR correction, p = 0.001-0.005). Hierarchical clustering by Bray-Curtis distance near-perfectly separated communities between PMA-treated and mock-treated samples, regardless of starting community ratios (Figure 4F, rug plot). Taxonomy analysis shows all mock-treatment communities had high relative abundance of *Bacillus* at 16 hours, an organism with a low relative abundance in the 16-hour PMA-treated samples (Figure 4F). This was not observed at the 72- and 196-hour time points (Supplemental Figure 4C-D). This suggests that many of the *Bacillus* reads detected in the mock-treatment samples before 24 hours could come from nonviable cells (presumably either spores or dead cells), while after 24 hours this is no longer the case.

**Figure 4.**
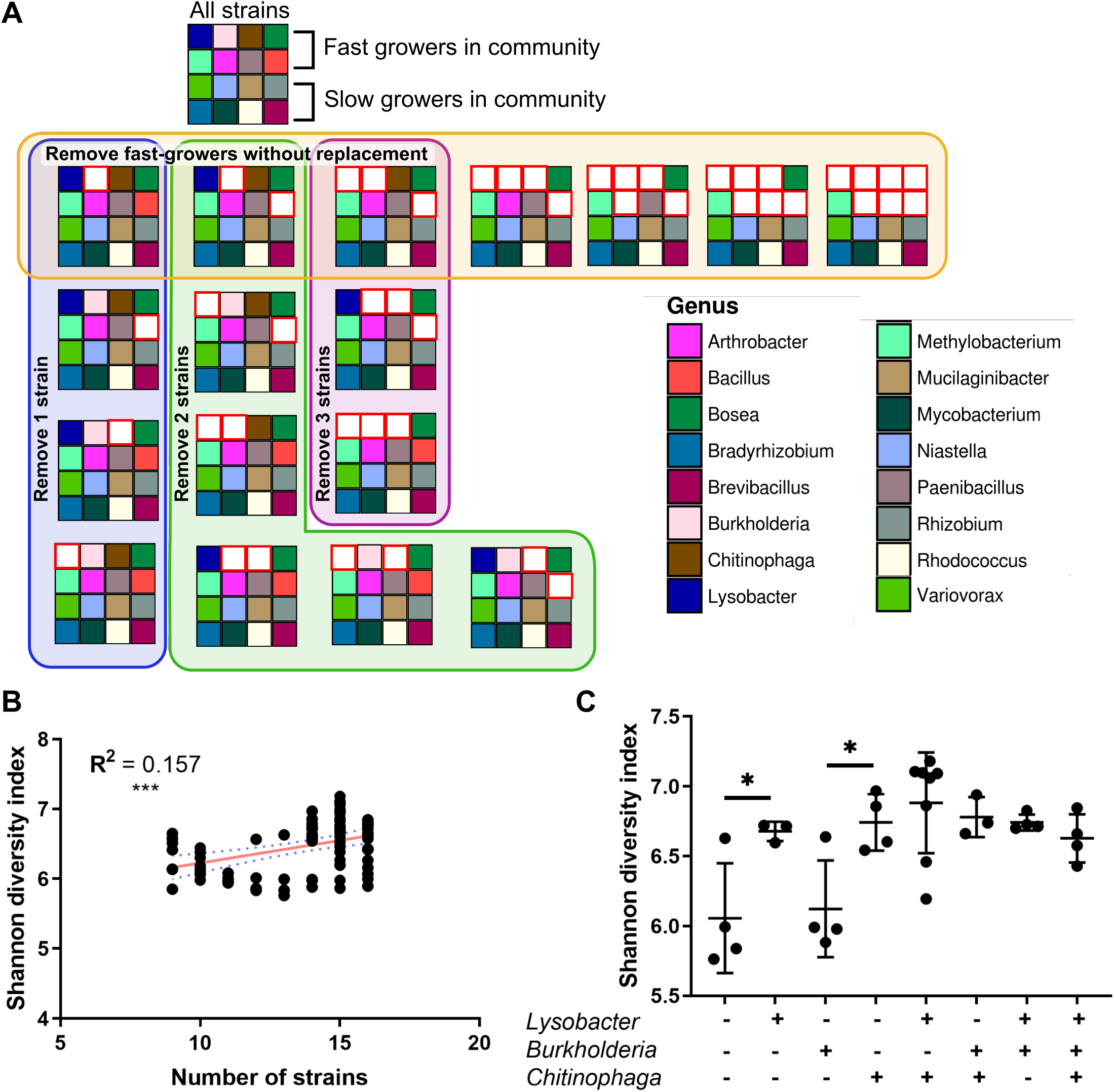
A) Schematic of 18 tested combinations of the 3x cutoff community, with each large square representing a different combination. Each small colored square represents an individual strain (see bottom right legend). White squares with a red border indicate the organism in that position was not included in that combination. For this experiment, strains were divided into fast-growing and slow-growing strains as indicated. *Sphingomonas* was not included in this experiment due to suspected contamination. **B)** The number of strains in each community combination plotted against the Shannon diversity index (n = 3-8 per combination). A linear regression trendline with 95% confidence interval is shown on the plot in red and blue, respectively. Spearman’s correlation coefficient is reported on the plot (*** p<0.001). **C)** Shannon diversity index for combinations with 13 or more strains. Communities are divided into groups based on the presence/absence of *Lysobacter, Burkholderia*, and *Chitinophaga* (* p<0.05).

### Community diversity dynamics are driven by presence of a few taxa

After determining that the highest α-diversity was observed in the 3x cutoff community composition, we sought to determine if the presence of specific taxa was required to generate this high-diversity community. For example, would there be a taxon or group of taxa whose removal caused community diversity to decrease sharply? To address this question, we started with the 3x cutoff composition and removed combinations of one or more organisms from the starting community. The absolute number of drops for the remaining organisms was left the same as before. We tested a total of 18 combinations within the 3x cutoff community (Figure 4A). We compared composition of the community with all 17 strains to the community with only “fast-growing” strains (strains that received 2 drops during community assembly), only “slow-growing” strains (received 2000 drops), and combinations that removed 1, 2, or 3 strains at a time, or removed the fast-growing strains one-by-one without replacement. Growth curves for each community are shown in Supplemental Figure 5.

**Figure 5.**
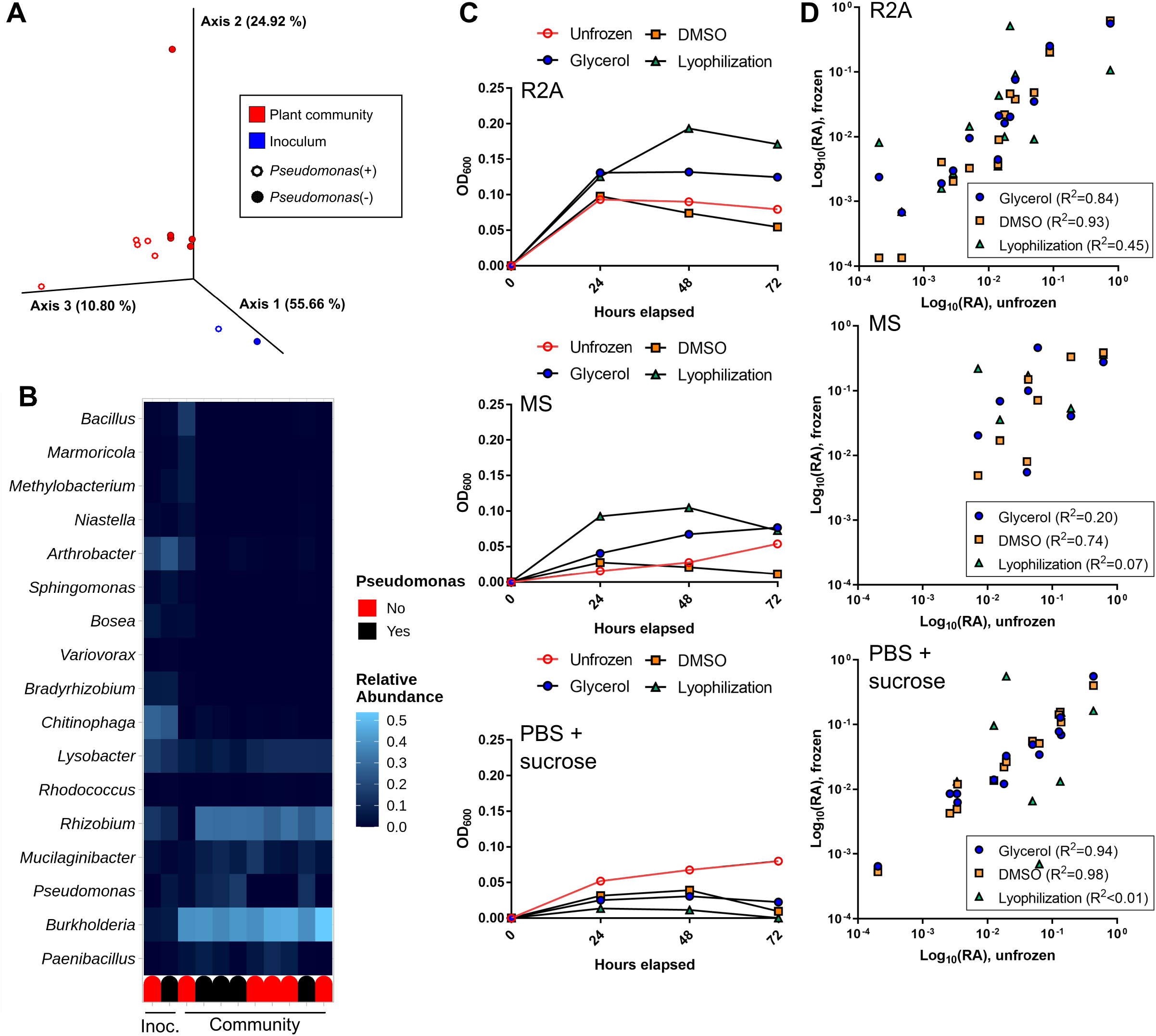
Community growth and composition with plant colonization and following cryopreservation. A-B) An equally-mixed community was inoculated to 12-day old *Brachypodium* plants, with or without *Pseudomonas*, and allowed to grow for 7 days. **A)** PCoA of Bray-Curtis distance between rhizosphere communities grown on plants for 7 days (n=5 each) and original inoculant (n=1). **B)** Heatmap of relative abundance of starting inoculum (Inoc.) and rhizosphere communities grown on plants for 7 days. Rug plot indicates presence (black) or absence (red) of *Pseudomonas* in inoculant. **C-D)** An equally-mixed community was preserved with 20% glycerol, 20% DMSO, or lyophilization and re-grown in R2A media (left panel), MS media (middle panel), or PBS with 10% sucrose (right panel). Growth and composition was compared to a community that was not frozen. **C)** Community growth measured through OD_600_. The unfrozen control community is shown in red. (n = 1) **D)** Comparison of log_10_(relative abundance) from 16S sequencing between frozen and unfrozen communities for each cryopreservation method. Each point represents the log_10_(relative abundance) of an individual genus in the frozen vs unfrozen community. Pearson’s correlation coefficient is reported for each comparison.

We first examined the effect of the total number of strains on community α-diversity, measured by Shannon diversity index (Figure 4B). There was a significant but very weak positive correlation between the number of strains in the starting community and final α-diversity (Spearman’s correlation coefficient R^2^=0.157, p=0.002). However, we were most interested in community combinations that deviated from the linear regression trendline shown in Figure 4B. We therefore analyzed the composition of the communities with 13 and more organisms for patterns that could explain large differences in α-diversity between communities. Each of these communities contained a “base community” of 14 organisms, with additional combinations of the fast-growing *Lysobacter*, *Burkholderia*, and *Chitinophaga* strains (Figure 4C, x-axis).

α-diversity varied significantly depending on the combination of 4 organisms present in the community. The addition of *Lysobacter* or *Chitinophaga* to the base community increased α-diversity, but the addition of *Burkholderia* did not. However, adding both *Lysobacter* and *Chitinophaga* did not significantly increase diversity over adding one of those organisms. Additionally, adding *Burkholderia* with *Lysobacter* or *Chitinophaga* did not reduce diversity. These results indicate that community α-diversity is not a simple additive effect of the individual organisms.

### Synthetic community is able to colonize the rhizosphere in EcoFAB system

To further our goal of developing a template for a model rhizosphere microbial community, we integrated our community with the EcoFAB device (https://eco-fab.org/), an existing system developed for reproducible studies with plants^16,18,37^. Colonization of plants in EcoFAB devices by the rhizosphere isolates would show these communities can be easily transferred to a current plant-microbiome system. To investigate this, sterile *Brachypodium distachyon* Bd21-3 seedlings were transferred into the EcoFAB device at 3 days after germination. 12-day-old plants were then inoculated with an equally-mixed community, either with or without *Pseudomonas*, and allowed to grow for 7 days (n=4-5). Rhizosphere community composition was then assessed with 16S sequencing and compared to the original inoculant.

Synthetic communities grown on plants were significantly different from the original inoculant, as determined by Bray-Curtis distance (Figure 5A; pairwise PERMANOVA with Benjamini-Hochberg FDR correction, p = 0.036). Communities with *Pseudomonas* were not significantly different from communities without *Pseudomonas*. However, a heatmap of relative abundance shows obvious changes in community composition between the inoculant and rhizosphere communities collected after 7 days on the plant (Figure 5B). *Burkholderia, Rhizobium*, and *Mucilaginibacter* increased in relative abundance, while several other organisms decreased in relative abundance. The increase in *Burkholderia* mirrors the presence of *Burkholderia* in the *in vitro* synthetic communities.

### Cryopreservation allows for community re-growth that recapitulates original community composition

To enable collaborative and comparable microbiome research, any synthetic community must grow and act reproducibly between different researchers and research institutions. We sought to determine which method of cryopreservation would best preserve community fidelity to an unfrozen community, to facilitate distribution of the synthetic rhizosphere community presented here to other research groups. We tested the growth and composition of an equally-mixed community following three methods of cryopreservation, compared to an unfrozen control community. The unfrozen community was allowed to grow for 72 hours in three types of media (R2A media, MS media, or PBS with 10% sucrose [the lyophilization medium]). Our cryopreserved communities were frozen at −80 °C for 3 days after lyophilization, or with either 20% glycerol or 20% DMSO as a cryopreservant. After freezing, the communities were subsequently grown for 3 days in the same media as the unfrozen community.

Community growth was highest in R2A media and lowest in PBS with 10% sucrose (Figure 5C). Comparison on log-transformed relative abundance values from 16S sequencing shows large differences in community composition between cryopreservation methods (Figure 5D). Lyophilization consistently produced the lowest Pearson’s coefficient of determination (R^2^ value) between frozen and unfrozen communities in all media. Glycerol and DMSO showed similar high coefficients in R2A and PBS with 10% sucrose media, although the DMSO coefficient was much higher than glycerol in MS media (R^2^=0.74, compared to R^2^=0.20).

## Discussion

In this study, we sought to develop a method to assemble and manipulate a synthetic soil community, while maintaining high levels of community α-diversity. Our results demonstrate the advantages of using a liquid-handling robot system to prepare synthetic communities. Direct comparison of hand-assembled and robot-assembled communities showed that machine-assembled communities have a generally lower level of dissimilarity than communities assembled by hand (Figure 2B). Examining community α-diversity (Supp Fig 1C), two of the four hand-assembly subjects produced communities with a similar standard deviation to the machine-assembled communities. However, the other two subjects produced communities with a larger standard deviation, indicating the differences inherent between hand-assembly subjects. Machine-assembly eliminates this source of variability in community production.

Other studies starting with a large number of community members^21,30^ reported a loss of detection of many organisms after the community was applied to plants. Of our 17 starting organisms (excluding *Pseudomonas)*, 11 were found consistently throughout all experiments and timepoints. *Pseudomonas simiae* was excluded from our later community experiments due to its tendency to proliferate rapidly in the community and decrease overall α-diversity. Additionally, community α-diversity was substantially increased by adjustment of starting organism ratios based on the growth rates of community members. Starting ratios containing orders of magnitude more of slower-growing organisms (2x cutoff and 3x cutoff) resulted in higher Shannon diversity index values than the 4 equally-mixed conditions (Figure 3B). The increases in α-diversity were seen even after 6 days of community growth. This indicates that synthetic community diversity can be increased over the diversity seen in typical 1:1 ratio communities by determining the growth rate of individual members and adjusting the starting ratios accordingly. As presented in this study, these ratios can also be determined through calculation of relative abundance changes during growth of an equally-mixed community.

In our 3x cutoff community without *Pseudomonas*, which displayed the highest α-diversity of the tested starting compositions, we further investigated the specific combination of organisms driving community diversity. We did not see a strong relationship between α-diversity and the total number of organisms in the starting community (Figure 4B). However, in communities with 14 or more organisms we noticed a range of α-diversity. We therefore analyzed the changes in α-diversity with the presence or absence of 3 specific fast-growing organisms in the community (*Bukholderia, Lysobacter*, and *Chitinophaga*). Our results indicate that different combinations of these organisms within the 3x cutoff community produce surprising non-linear diversity results. The addition of *Lysobacter* or *Chitinophaga* to the 13-member base community resulted in significant increases in α-diversity (Figure 4C), while the addition of *Burkholderia* did not change the α-diversity. Adding both *Lysobacter* and *Chitinophaga* to the base community resulted in a slight increase in diversity in most replicates, although this did not reach significance. However, adding *Lysobacter, Chitinophaga*, and *Burkholderia* together reduced diversity to levels similar to *Lysobacter* or *Chitinophaga* alone. These results show that community α-diversity is not driven solely by additive effects of individual community members. Interactions between organisms can alter the effect of individual microbes on diversity. How these interactions move beyond diversity to affect individual organism metabolism is still an open question.

To address the potential influence of relic DNA on our sequencing results, we tested the effect of PMA treatment on the community ratios with highest α-diversity. Our results indicate that relic DNA can have a significant effect on sequencing results from <24 hours of community growth. Significantly fewer quality sequencing reads were detected in PMA-treated communities after 0, 16, and 72 hours of growth. However, the taxonomy relative abundance values were similar from 72 hours out to 196 hours (8 days). This indicates that while relic DNA can significantly alter sequencing results in our system in short-term growth studies, this effect diminishes at later time points (between 24 and 72 hours).

After investigating and optimizing α-diversity for our synthetic rhizosphere community, we next sought to display the utility of this community in a controlled model microbiome system. We tested community colonization of the EcoFAB device, a system designed for reproducible plant-microbiome system studies. The synthetic community was able to colonize the rhizosphere of *Brachypodium distachyon* plants grown in the EcoFAB device and was significantly different from the original inoculant after 7 days growth on the plant (Figure 5A-B). The presence of *Pseudomonas* did not significantly change the community in the EcoFAB system, unlike what was seen in our *in vitro* synthetic community system. The high relative abundance of *Burkholderia* in the rhizosphere communities was similar to levels of *Burkholderia* seen in the *in vitro* community, while other organisms had different relative abundance levels. These differences were expected given the addition of the *Brachypodium* plant, which produces factors that affect soil microbe growth and metabolism. Indeed, aspects of plant-associated microbiomes have been shown to change rapidly in the natural environment based on climatic factors^41^. Future studies will focus on improving the accuracy of the *in vitro* by adding plant factors to the growth media.

We additionally determined the optimal method for community cryopreservation and re-growth. Although sequencing results indicated a high correlation between the glycerol and DMSO frozen communities and unfrozen community in PBS with 10% sucrose, the low OD_600_ values in this medium indicates this fidelity is likely due to a lack of growth following thawing. Preservation of 20% glycerol and re-growth in MS media led to similar growth by OD_600_ as the unfrozen community, but sequencing results revealed a poor correlation. However, glycerol preservation and re-growth in 0.1X R2A media resulted in community re-growth that closely re-capitulated the unfrozen community in both OD_600_ and sequencing results.

Community reproducibility, diversity, and preservation are essential questions to be addressed in the development of reproducible microbiome model communities. Developing a defined and reproducible synthetic microbial community, accounting for various starting organism ratios, and the ability to preserve communities for dissemination are key elements to aid reproducible microbiome sciences. Additionally, we have shown that our synthetic community can be used in EcoFAB devices to reproducibly study plant-microbe interactions in the rhizosphere. The methods and workflows developed here can be readily adapted for the design and study of other model communities and to standardize microbiome research.

## Materials and Methods

### Isolate selection

Isolates were selected from a collection obtained from the rhizosphere and soil surrounding a single switchgrass plant grown in marginal soils described elsewhere^42,43^. Isolates are available from the Leibniz Institute German Collection of Microorganisms and Cell Cultures GmbH (DSMZ) under accession numbers DSM 113524, DSM 113525, DSM 113526, and DSM 113527 (remaining numbers will be included prior to publication).

### Soil isolate growth conditions

Individual isolates were grown in 3-5mL liquid cultures of 1X R2A media (Teknova, cat # R0005) in 14-mL culture tubes in aerobic conditions, 30 °C, without shaking. Isolates were allowed to grow for 5-7 days before diluting for community generation. 0.2X and 0.1X media was made by diluting 1X media with water purified by a Milli-Q water purification system and vacuum-filtering through a 0.22μM filter. Growth curves for individual isolates were conducted in 96-well plates.

Isolates cultured in 1X R2A media were diluted to a starting OD_600_ of 0.05 in 200 uL of 0.1x R2A. Sterile R2A media served as the negative control.

### Synthetic community growth conditions

Communities were grown in 200μL of liquid R2A media in 96-well plates in aerobic conditions, 30 °C, without shaking. To prevent condensation, each plate lid was coated with 3mL of an aqueous solution with 20% ethanol and 0.01% Triton X-100 (Sigma, cat # X100-100ML). Excess liquid was removed after 30 seconds and the lid was allowed to air-dry for 30min under a UV light for sterilization. To further prevent condensation, plates were set on 4 100mm-diameter Petri dishes (2 stacks of 2 dishes) filled with ~20mL water each to generate a humid environment around the plates. Optical density readings at 600nm, to normalize isolates and monitor community growth, were taken with a Molecular Devices SpectraMax M3 Multi-Mode Microplate Reader (VWR, cat # 89429-536).

### Synthetic community assembly using the CellenONE printer

Communities were assembled using a SCIENION CellenONE machine (https://www.scienion.com/). Individual isolate cultures were OD_600_-normalized to 0.025 (after subtracting media blank), then transferred from a 384- or 96-well probe plate to a 96-well target plate using a CellenONE piezo dispense capillary (PDC) (size medium; Scienion, cat # P-20-CM) with droplet size set to 390-420 picoliters. The number of drops per isolate for each community was programmed by hand using the provided Scienion software (v1.92). The number of drops per organism for each ratio can be found in Supplemental Table 3. Droplet integrity was confirmed before and after each isolate spotting run using the droplet camera and automated droplet detection. The PDC was cleaned between isolates by flushing the PDC interior with 0.5mL water. 200 drops of R2A were added to negative control wells as the last step in each experimental setup, to ensure no contamination occurred due to incomplete flushing of the PDC between isolates. For the community dynamics experiment, organisms receiving 2000 drops were added to communities with a multichannel pipettor.

### Treatment with PMA to remove relic DNA

PMA (Biotium, cat # 40013) was added to communities to a final concentration of 10μM directly prior to sample collection (PMA-treatment); 5μL water (mock-treatment) was added to control communities. Communities were then incubated in the dark for 5 minutes at room temperature, then placed <15cm from a direct fluorescent light source and incubated on ice for 30min. Communities were then frozen at −20 °C until processing for sequencing.

### Plant colonization experiment

*Brachypodium distachyon* Bd21-3 seeds were dehusked, sterilized, and germinated on 0.1X Murashige and Skoog (MS) basal salt mixture M524 plates, pH 5.7 (Phyto Technology Laboratories) in a 250 μmol/m^2^ s-1 16-hr light/8-hr dark regime at 24 °C for three days. EcoFABs were sterilized as previously described^18^, and seedlings transferred to EcoFAB chambers filled with 0.1X MS at 3 days after germination. Twelve-day old plants were inoculated with an equally-mixed community of 17 or 18 bacterial isolates, as described above. To mix the community, the OD_600_ of each isolate was measured, with the assumption that OD_600_ of 1 is equal to ~10^9 CFU (colony-forming units)/mL^31^. Isolates were combined at 10^5 CFU/mL per isolate in the final EcoFAB volume. Plants were harvested seven days after inoculation. Microbial communities were detached from the plant root by vortexing the root in 0.1 phosphate-buffered saline (PBS) for 10 minutes at maximum speed, followed by centrifugation at 10000g, at 6 °C. DNA was extracted by using the Qiagen DNeasy PowerSoil Pro kit according to manufacturer’s instructions (cat # 47014).

### Community cryopreservation and re-growth

All community members were OD_600_-normalized to 0.1 after 3 days of growth in 1X R2A and mixed equally to a final estimated total CFU count of 7.2*10^8 CFU. The community was then centrifuged (5000g, 5min) and resuspended in 4mL of 0.1X R2A media. 250μL of the community was inoculated into 4mL of 0.1X R2A, MS media (RPI, cat # M10200), or PBS + 10% sucrose (w/v) as the “unfrozen” control community. 500μL of the community was mixed with 500μL of either 40% glycerol, 40% DMSO, or PBS with 20% sucrose (w/v). The glycerol and DMSO stocks were frozen immediately at −80 °C. The PBS with 10% sucrose stock was lyophilized on a Labconco FreeZone Plus Freeze Dry System (cat # 7386030) and then stored at −80 °C. Stocks were thawed after 3 days and 250μL of stock was inoculated into the same 3 types of media as the unfrozen community. Samples from all communities were frozen at −20 °C after 3 days of growth for 16S sequencing analysis.

### DNA extraction and sequencing

DNA extracted with a kit was processed with the Qiagen DNeasy PowerSoil Pro kit according to manufacturer’s instructions (cat # 47014). DNA extracted by boiling was processed by thawing community samples, transferring 100μL to a PCR plate, and heating the plate in a PCR machine at 100 °C for 10 minutes. 5μL of undiluted sample was used as DNA input for the 16S rRNA gene amplicon library protocol. 16S libraries for the cryopreservation, adjusted community ratios, PMA, and boil-extraction comparison experiments were prepared using 515F-806R primers according to the Earth Microbiome Project protocol^44^ and sequenced on an Illumina MiSeq platform with a paired-end 150 V2 kit as previously described^45,46^. 16S libraries for the community dynamics experiment were prepared using 341F-805R primers (F 5’-CCTACGGGNGGCWGCAG-3’ R 5’-GACTACHVGGGTATCTAATCC-3’) and sequenced on an Illumina MiSeq platform with a paired-end 150 V2 kit. 16S libraries for the plant experiments were prepared using 515F-806R primers and sequenced on an Illumina NovaSeq platform with a paired-end 250 V2 kit. Shotgun metagenomics libraries for the human-/machine-assembled experiment were prepared using 1ng DNA input and Nextera XT indexes and sequenced on an Illumina MiSeq platform with a paired-end 150 V2 kit. DNA sequences generated through this study are available on the NCBI Sequence Read Archive (submission SUB9617801).

### 16S rRNA sequencing analysis and statistical analyses

All 16S sequences were analyzed using QIIME2^47^ (v2020-11). Paired-end reads were joined using the “qiime vsearch join-pairs” command and quality-filtered and denoised (using default parameters) with Deblur^48^. Reads were trimmed as appropriate for quality for each experiment (150bp for human-/machine-assembled, cryopreservation, adjusted community ratios, and PMA experiments; 200bp for plant experiment). α- and β-diversity was calculated using the “qiime diversity” set of commands, with alpha rarefaction used to determine an appropriate sampling depth. Robust Aitchison distance was calculated using the DEICODE plugin^49^. Microbial taxonomy was assigned to the filtered sequences with the “qiime feature-classifier classify-sklearn” command, using a scikit-learn classifier created from a custom database of the 16S rRNA gene sequences for the isolates used in the study. Heatmaps and relative abundance plots were generated using R^50^ (v3.3.2) with the packages dplyr^51^, phyloseq^52^, ggplot2^53^, and scales^54^f. β-diversity plots were generated using QIIME2. All other plots were generated using GraphPad Prism 7 software. All code used to process and analyze sequencing results can be accessed through Github at https://github.com/jkccoker/Soil_synthetic_community.

### Shotgun metagenomics sequencing analysis

Shotgun sequencing data were quality-filtered during adapter trimming with Trimmomatic^55^ (v0.36) using the settings “ILLUMINACLIP:NexteraPE-PE.fa:2:30:10 LEADING:10 TRAILING:10 SLIDINGWINDOW:4:15 MINLEN:36”. Trimmed reads were aligned to a custom database of community strain genomes using bowtie2^56^ (v2.2.3) using default settings. α- and β-diversity was calculated in phyloseq. β-diversity plots were generated using phyloseq. All other plots were generated using GraphPad Prism 7 software.

## Acknowledgements

The development of the technologies and research described in this article were funded through Trial Ecosystem Advancement for Microbiome Science Program and the Microbial Community Analysis and Functional Evaluation in Soils (m-CAFEs) Science Focus Area Program at Lawrence Berkeley National Laboratory funded by the U.S. Department of Energy, Office of Science, Office of Biological & Environmental Research Awards DE-AC02-05CH11231. This material is also based upon work supported by the U.S. Department of Energy, Office of Science, Office of Biological & Environmental Research under Awards DE-SC0021234 and DE-SC0022137.

---

**Supplemental Figure 1.** Comparison of boiling and conventional kit DNA extraction for rhizosphere isolates (A-B). Isolates were mixed in equal amounts, as determined by OD normalization. DNA was extracted either by heating to 95°C for 10 minutes in a PCR machine (boiling) or using a conventional extraction kit (Qiagen PowerSoil Pro) (extraction). (n=3 per condition) **A)** Taxonomy of samples through 16S sequencing. **B)** Comparison of a logarithmic transformation of mean relative abundance values in boiling and extraction samples. Pearson’s correlation coefficient is shown on the plot. Community diversity between assembly methods and media dilutions (C-D). **C)** Shannon diversity index of machine-assembled and human-assembled communities from 4 different people (n=4-8 each). **D)** Observed operational taxonomic units (OTUs) for equally-mixed community grown in 1X, 0.2X, and 0.1X R2A media for 3 days (n=8 each).

**Supplemental Figure 2.** *Growth curves of individual rhizosphere isolates in 0.1X R2A media*.

**Supplemental Figure 3.** Growth curves of communities with adjusted starting ratios. Community growth was monitored through OD600 717 for 6 days. **A)** Growth of communities without Pseudomonas simiae. **B)** Growth of communities with Pseudomonas simiae.

**Supplemental Figure 4.** Growth curves and taxonomic composition of PMA-treated and mock-treated communities. Community growth was monitored through OD600 for 8 days (n=4 for each condition). **A)** Growth of communities that were not treated with PMA before sample collection **B)** Growth of communities treated with PMA before sample collection. **C)** Heatmap of taxonomic composition after 72 hours of growth, all community ratios combined. **D)** Heatmap of taxonomic composition after 196 hours (8 days) of growth, all community ratios combined. PMA-treated (yellow) and mock-treated (red) communities are marked in the rug plot at the bottom of each heatmap.

## Notes

https://www.ncbi.nlm.nih.gov/bioproject/PRJNA807292/

https://github.com/jkccoker/Soil_synthetic_community

### Competing Interest Statement

The authors have declared no competing interest.

