## Supplemental figures for "A reproducible and tunable synthetic soil microbial community provides new insights into microbial ecology"

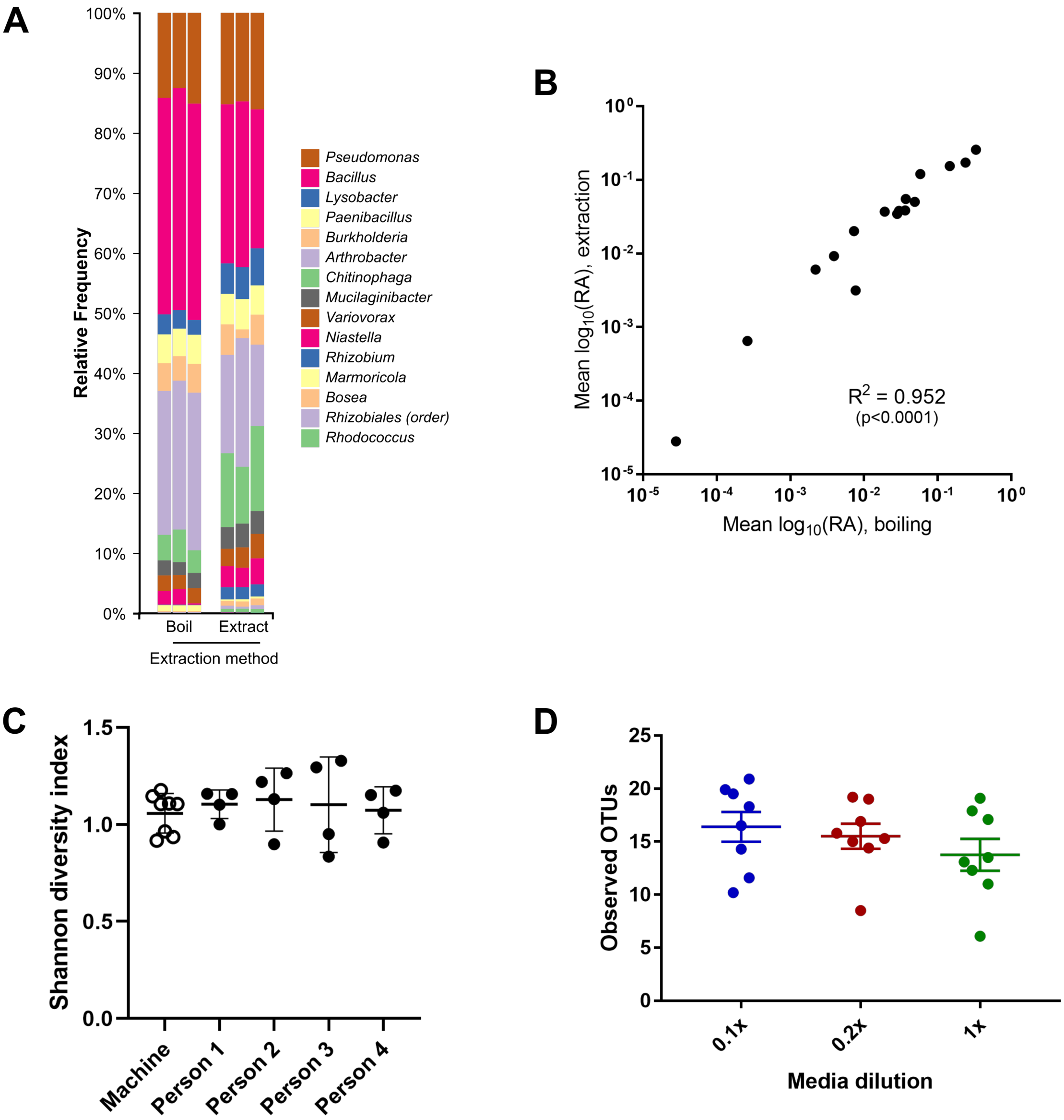


**Supplemental Figure 1.** *Comparison of boiling and conventional kit DNA extraction for rhizosphere isolates (A-B).* Isolates were mixed in equal amounts, as determined by OD normalization. DNA was extracted either by heating to 95°C for 10 minutes in a PCR machine (boiling) or using a conventional extraction kit (Qiagen PowerSoil Pro) (extraction). (n=3 per condition) **A)** Taxonomy of samples through 16S sequencing. **B)** Comparison of a logarithmic transformation of mean relative abundance values in boiling and extraction samples. Pearson's correlation coefficient is shown on the plot. *Community diversity between assembly methods and media dilutions (C-D).* **C)** Shannon diversity index of machine-assembled and human-assembled communities from 4 different people (n=4-8 each). **D)** Observed operational taxonomic units (OTUs) for equally-mixed community grown in 1X, 0.2X, and 0.1X R2A media for 3 days (n=8 each).

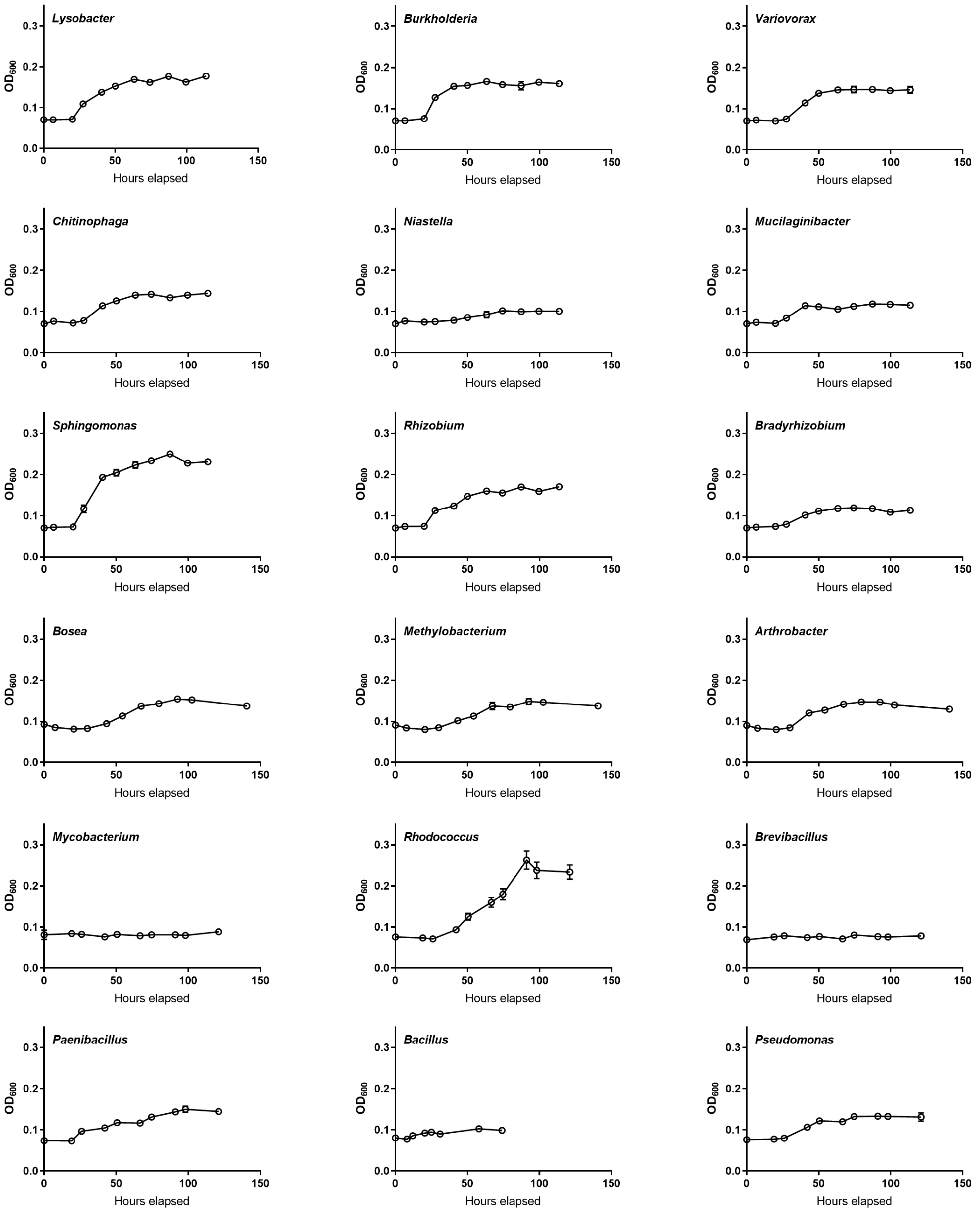


**Supplemental Figure 2.** *Growth curves of individual rhizosphere isolates in 0.1X R2A media.*

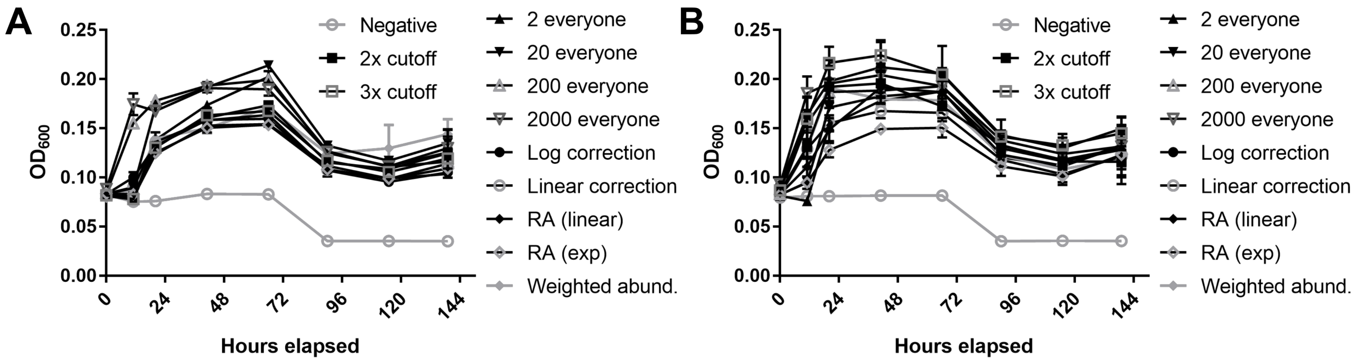


**Supplemental Figure 3.** *Growth curves of communities with adjusted starting ratios*. Community growth was monitored through $OD_{600}$ for 6 days. **A)** Growth of communities without *Pseudomonas simiae*. **B)** Growth of communities with *Pseudomonas simiae*.

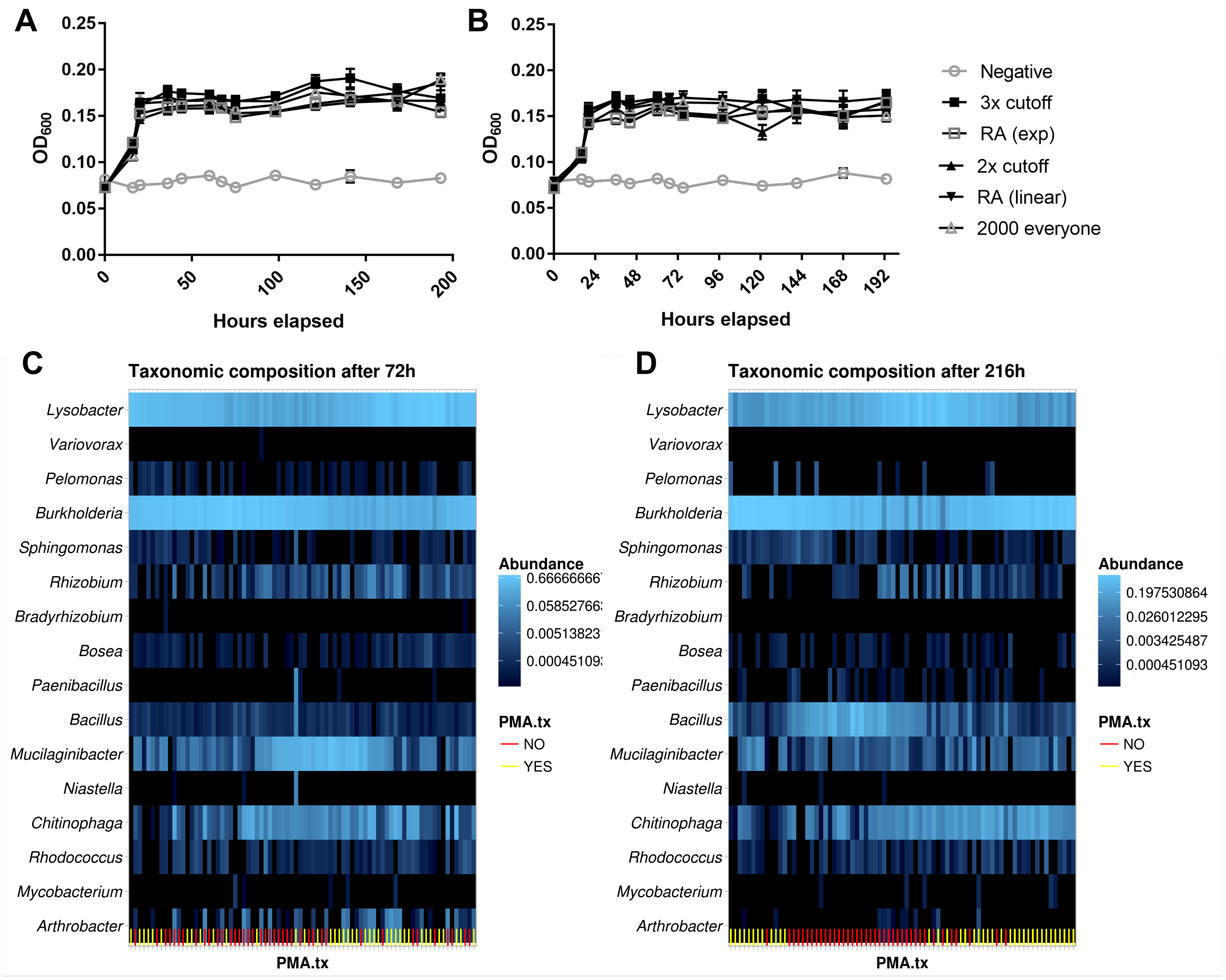


**Supplemental Figure 4.** *Growth curves and taxonomic composition of PMA-treated and mock-treated communities*. Community growth was monitored through $OD_{600}$ for 8 days (n=4 for each condition). **A)** Growth of communities that were not treated with PMA before sample collection **B)** Growth of communities treated with PMA before sample collection. **C)** Heatmap of taxonomic composition after 72 hours of growth, all community ratios combined. **D)** Heatmap of taxonomic composition after 196 hours (8 days) of growth, all community ratios combined. PMA-treated (yellow) and mock-treated (red) communities are marked in the rug plot at the bottom of each heatmap.
