## Supplemental tables for "A reproducible and tunable synthetic soil microbial community provides new insights into microbial ecology"

**Supplemental Table 1.** *Strain isolates used in this study and examples of plants associated with them in previous publications.*

| **Genus** | **Strain** | **Associated plants**[a] |
|---|---|---|
| *Lysobacter* | OAE881 | Nicotiana tabacum L, tomato, pepper[1–3] |
| *Burkholderia* | OAS925 | Zea mays L, Betula, Equisetum, Quercus, Senecio vulgaris, Triticum aestivum, Zantedeschia, Coffea, Saccharum officinarum, Zea mays L, Zea mays L, Lolium multiflorum, citrus, wheat[4–9] |
| *Variovorax* | OAS795 | Citrus, maize, tomato, wheat[8,10,11] |
| *Chitinophaga* | OAE865 | Tomato, Oryza sativa L, Cymbidium goeringii, ginseng[12–15] |
| *Niastella* | OAS944 | Hibiscus syriacus L, persimmon tree, Populus euphratica, Korean ginseng[16–19] |
| *Mucilaginibacter* | OAE612 | Gossypium hirsutum L, Angelica sinensis, ginseng, Dokdo Island (S Korea)[20–23] |
| *Sphingomonas* | OAE905 | Citrus, wheat, Oryza sativa L[8–10,24,25] |
| *Rhizobium* | OAE497 | Citrus, Oryza sativa, Dioscorea alata, Dioscorea esculenta[10,26,27] |
| *Bradyrhizobium* | OAE829 | Citrus, Vaccinium angustifolium, wheat, Brazilian sugarcane[8,10,28,29] |
| *Bosea* | OAE506 | Zea mays L, Cyperus rotundus L[30,31] |
| *Methylobacterium* | OAE516 | Eucalyptus spp., Oryza sativa cv. Dongjin and Lycopersicon esculentum L. cv. Mairoku, maize[32–34] |
| *Arthrobacter* | OAP107 | Ginkgo biloba L, Quercus ilex, Triticum aestivum L (wheat)[35–37] |
| *Mycobacterium* | OAE908 | Soil from Haikou (China), tomato, Oryza sativa L. cv. Wusimi[38–40] |
| *Rhodococcus* | OAS809 | Zea mays L, Oryza sativa L.[30,41] |
| *Brevibacillus* | OAP136 | Zea mays L, Lolium perenne, Pinellia ternata, Nicotiana tabacum L, Gossypium hirsutum[30,42–45] |
| *Paenibacillus* | OAE614 | Solanum lycopersicum, Oryza sativa L, wheat[46–48] |
| *Bacillus* | OAE603 | Zea mays L, Triticum aestivum L (wheat), Lolium perenne, Nicotiana tabacum L[30,37,42,44] |
| *Pseudomonas simiae* | WCS417 | Wheat, citrus, Zea mays L, Nicotiana tabacum L[9,10,30,44] |

[a]References were found through PubMed searches (conducted on 8/17/2021) of “<genus> rhizosphere”, “<genus> soil isolation”, and/or “<genus> plant isolation”. Plant scientific names are listed when included in the references; otherwise plant common names are used.

**Supplemental Table 2.** *Relative abundance values of organisms from an equally-mixed community.*

| | SRA[a] | FRA[b] | F/S ratio (FRA/SRA)[c] |
|---|---|---|---|
| *Lysobacter* OAE881 | 0.029507 | 0.160573 | 5.44179057 |
| *Pseudomonas simiae* WCS417 | 0.139359 | 0.570065 | 4.09063447 |
| *Sphingomonas* OAE905 | 0.000018 | 0.000053 | 2.8754562 |
| *Burkholderia* OAS925 | 0.044979 | 0.117325 | 2.60846168 |
| *Rhizobium* OAE497 | 0.001012 | 0.001477 | 1.45909576 |
| *Bacillus* OAE603 | 0.363800 | 0.137535 | 0.37805086 |
| *Chitinophaga* OAE865 | 0.044788 | 0.007738 | 0.17275925 |
| *Mucilaginibacter* OAE612 | 0.023927 | 0.002400 | 0.10030911 |
| *Bosea* OAE506 | 0.001536 | 0.000085 | 0.0554598 |
| *Rhodococcus* OAS809 | 0.000219 | 0.000012 | 0.05523912 |
| *Paenibacillus* OAE614 | 0.047480 | 0.002502 | 0.05269173 |
| *Niastella* OAS944 | 0.016974 | 0.000045 | 0.0026626 |
| *Variovorax* OAS795 | 0.024828 | 0.000059 | 0.00237307 |
| *Arthrobacter* OAP107 | 0.250202 | 0.000049 | 0.0001968 |
| *Bradyrhizobium* OAE829 | 0.001405 | 0.000000 | 0 |
| *Methylobacterium* OAE516 | 0.001405 | 0.000000 | 0 |
| *Mycobacterium* OAE908 | 0.000120 | 0.000000 | 0 |
| *Brevibacillus* OAP136 | 0.000000 | 0.000019 | 0 |

[a]SRA = starting relative abundance; RA reported by 16S sequencing at time 0 of an equally-mixed community.
[b]FRA = final relative abundance; RA reported by 16S sequencing after 3 days growth of an equally-mixed community.
[c]F/S ratio = fold-change in relative abundance between time 0 and 3 days, calculated as FRA / SRA

**Supplemental Table 3.** *Equations for starting community ratios and number of CellenONE printer drops per organism.*

| | 2 everyone | 20 everyone | 200 everyone | 2000 everyone | 2x cutoff | 3x cutoff | Linear correction | Log correction | RA (exp) | RA (linear) | Weighted abundance |
|---|---|---|---|---|---|---|---|---|---|---|---|
| **Equation[a]** | 2 drops | 20 drops | 200 drops | 2000 drops | 2 or 2000 drops | 2, 200, or 2000 drops | Drops = (10-FSR)*10 | Drops = $(10-FSR)^{10}/2e6$ | Drops = $100/SRA^2$ | Drops = 1/SRA*10 | Drops = 100*2^((1-SRA)*(1-FSR)) |
| *Lysobacter* | 2 | 20 | 200 | 2000 | 2 | 2 | 46 | 2 | 2 | 3 | 200 |
| *Pseudomonas* | 2 | 20 | 200 | 2000 | 2 | 2 | 59 | 26 | 2 | 2 | 16 |
| *Sphingomonas* | 2 | 20 | 200 | 2000 | 2 | 2 | 71 | 168 | 2986 | 5464 | 27 |
| *Burkholderia* | 2 | 20 | 200 | 2000 | 2 | 2 | 74 | 243 | 2 | 2 | 34 |
| *Rhizobium* | 2 | 20 | 200 | 2000 | 2 | 2 | 85 | 1033 | 2 | 99 | 73 |
| *Bacillus* | 2 | 20 | 200 | 2000 | 2 | 200 | 96 | 3401 | 2 | 2 | 132 |
| *Chitinophaga* | 2 | 20 | 200 | 2000 | 2 | 200 | 98 | 4200 | 2 | 2 | 173 |
| *Mucilaginibacter* | 2 | 20 | 200 | 2000 | 2 | 200 | 99 | 4520 | 2 | 4 | 184 |
| *Bosea* | 2 | 20 | 200 | 2000 | 2 | 200 | 99 | 4730 | 2 | 65 | 192 |
| *Rhodococcus* | 2 | 20 | 200 | 2000 | 2 | 200 | 99 | 4731 | 21 | 457 | 192 |
| *Paenibacillus* | 2 | 20 | 200 | 2000 | 2 | 200 | 99 | 4743 | 2 | 2 | 187 |
| *Niastella* | 2 | 20 | 200 | 2000 | 2000 | 2000 | 100 | 4987 | 2 | 6 | 197 |
| *Variovorax* | 2 | 20 | 200 | 2000 | 2000 | 2000 | 100 | 4988 | 2 | 4 | 196 |
| *Arthrobacter* | 2 | 20 | 200 | 2000 | 2000 | 2000 | 100 | 4999 | 2 | 2 | 168 |
| *Bradyrhizobium* | 2 | 20 | 200 | 2000 | 2000 | 2000 | 100 | 5000 | 2 | 71 | 200 |
| *Methylo-bacterium* | 2 | 20 | 200 | 2000 | 2000 | 2000 | 100 | 5000 | 2 | 71 | 200 |

| | | | | | | | | | | | |
|---|---|---|---|---|---|---|---|---|---|---|---|
| *Mycobacterium* | 2 | 20 | 200 | 2000 | 2000 | 2000 | 100 | 5000 | 70 | 837 | 200 |
| *Brevibacillus* | 2 | 20 | 200 | 2000 | 2000 | 2000 | 100 | 5000 | 5739 | 7575 | 200 |

[a]FSR = F/S ratio, as defined in Supplemental Table 1. The minimum number of drops per organism was set at 2.
